# Bacterial T6SS Effector EvpP Inhibits Neutrophil Recruitment via Jnk-Caspy Inflammasome Signaling *In vivo*

**DOI:** 10.1101/453076

**Authors:** Jinchao Tan, Dahai Yang, Zhuang Wang, Xin Zheng, Yuanxing Zhang, Qin Liu

## Abstract

The type VI secretion system (T6SS) comprises dynamic complex bacterial contractile nanomachines and is used by many bacteria to inhibit or kill other prokaryotic or eukaryotic cells. Previous studies have revealed that T6SS is constitutively active in response to various stimuli, or fires effectors into host cells during infection. It has been proposed that the T6SS effector EvpP in *Edwardsiella piscicida* can inhibit NLRP3 inflammasome activation via the Ca^2+^-dependent JNK pathways. Here, we developed an *in vivo* infection model by microinjecting bacteria into the tail vein muscle of 3-day-post-fertilized zebrafish larvae, and found that both macrophages and neutrophils are essential for bacterial clearance. Further study revealed that EvpP plays a critical role in promoting the pathogenesis of *E. piscicida* via inhibiting the phosphorylation of Jnk signaling to reduce the expression of *cxcl8a*, *mmp13* and *IL-1β in vivo.* Subsequently, by utilizing *Tg* (*mpo:eGFP*^+/+^) zebrafish larvae for *E. piscicida* infection, we found that the EvpP-inhibited Jnk-caspy inflammasome signaling axis significantly suppressed the recruitment of neutrophils to infection sites, and the *caspy*‐ or *IL-1β*-MO knockdown larvae were more susceptible to infection and failed to restrict bacterial colonization *in vivo*.

**IMPORTANCE:** Innate immunity is regulated by phagocytic cells and is critical for host control of bacterial infection. In many bacteria, T6SSs can affect bacterial virulence in certain environments, but little is known about the mechanisms underlying T6SS regulation of innate immune responses during infection *in vivo*. Here, we investigated the role of an *E. piscicida* T6SS effector EvpP in manipulating the reaction of neutrophils *in vivo.* We show that EvpP inhibits the activation of Jnk-caspy inflammasome pathway in zebrafish larvae, and reveal that macrophages are essential for neutrophil recruitment *in vivo*. This interaction improves our understanding about the complex and contextual role of a bacterial T6SS effector in modulating the action of myeloid cells during infection, and offers new insights into the warfare between bacterial weapons and host immunological surveillance.

---

The bacterial type VI secretion system (T6SS) is a versatile secretion system capable of facilitating a variety of interactions with eukaryotic hosts and/or bacterial competitors (1). T6SS effectors are delivered upon cell-to-cell contact and include factors engaged in interbacterial competition and those that mediate pathogenicity in the context of eukaryotic host infections (2-5). To date, various anti-bacterial effectors have been identified that attack the bacterial cell wall, nucleases, or lipases via diverse activities, including those of muramidases and peptidases (1). There are several bacterial species that utilize the T6SS to mediate pathogenicity in eukaryotic hosts, including *Vibrio cholerae*, *Pseudomonas aeruginosa*, *Burkholderia pseudomallei*, and *Aeromonas hydrophila* in mammals, and *Edwardsiella piscicida* in fish. Although many molecular consequences of T6SS activity on eukaryotic cells have been deciphered (1), few anti-eukaryotic effectors have been identified other than the enzymatic domains found in VgrGs (2) and the phospholipase D (PLD) enzyme PldB (6). Recently, Chen *et al.* identified a non-VgrG T6SS effector, EvpP, from *E. piscicida*, and revealed its role in inhibiting NLRP3 inflammasome activation in macrophages. However, little is known about the physiological role of this bacterial T6SS effector in manipulating host immunity during pathogenic infection *in vivo*.

The zebrafish (*Danio rerio*) is a genetically and optical accessible model for infectious diseases (7-9), in which the *in vivo* innate immune responses can be studied in the context of a whole organism. Using zebrafish larvae, infectious processes can be described in detail using *in vivo* imaging techniques because of their small size and transparency during the first week after fertilization. Thus, zebrafish larvae have been used to analyze the innate immune response after bacterial infections, including *Mycobacterium marinum* (10), *Streptococcus* sp. (11), *Salmonella typhimurium* (12), *Staphylococcus aureus* (13), and *Burkholderia cenocepacia* (14). Moreover, zebrafish are increasingly used to study the function of neutrophils and host pathogen interactions, and the generation of transgenic zebrafish lines with fluorescently labeled leukocytes has made it possible to visualize neutrophil responses to infection in real time (15).

*Edwardsiella piscicida*, previously named as *E. tarda* (16), is an intracellular bacterium with broad cellular tropism; it can infect practically all vertebrates, causing septicemia and fatal infections (17, 18). T3SS and T6SS have been identified as important components of virulence in this pathogen (19-21). Moreover, *E. piscicida* activates NLRC4 and NLRP3 inflammasomes via T3SS and inhibits the NLRP3 inflammasome via EvpP (22). To date, although several infection models have been used to explore the biology of *Edwardsiella* sp. (23, 24), the events of myeloid cell responses during *E. piscicida* infection *in vivo* remain to be clarified. In this study, we established an microinjection infection model in the tail vein muscle of 3-day-post-fertilized zebrafish larvae and analyzed the role of T6SS effector, EvpP, in manipulating host immune responses *in vivo*. We demonstrated that EvpP inhibits the phosphorylation of JNK-MAPK pathway, subsequently suppressing the caspy-inflammasome signaling cascades, contributing to the inhibition of neutrophils recruitment. Moreover, we found that both macrophages and neutrophils are critical for the clearance of *E. piscicida in vivo*. Collectively, this study advances our understanding of the mechanisms of the bacterial T6SS effector in regulating the action of innate immune cells during infection.

## RESULTS

### Macrophages and neutrophils are critical for *E. piscicida* infection

To analyze the functional roles of *E. piscicida* during infection *in vivo*, we determined a reproducible route of infection with rapid kinetics by tail muscle microinjection of 3-day-post-fertilized (dpf) larvae with the indicated doses of *E. piscicida* (Fig. 1A). As shown in Fig. 1B, *E. piscicida* microinjection-infection induced mortality in a dose-dependent manner. Mortality began at 24 h postinfection (hpi), and consistently reached 100% between 24 to 48 hpi when infected with 100 cfu/larvae. Nearly 60% of larvae succumbed when infected with 50 cfu/larvae, and ~20% larvae succumbed when infected with 10 cfu/larvae. Based on these results, we calculated the LD_50_ of wild type *E. piscicida* as 45 cfu/larvae. To further visualize *E. piscicida* infection *in vivo*, we constructed mCherry-labeled *E. piscicida* strains to microinject into the tail muscle, and found that *E. piscicida* colonized at the infection site and diffused from 12 to 24 hpi (Fig. 1 C and D).

**FIG 1.**
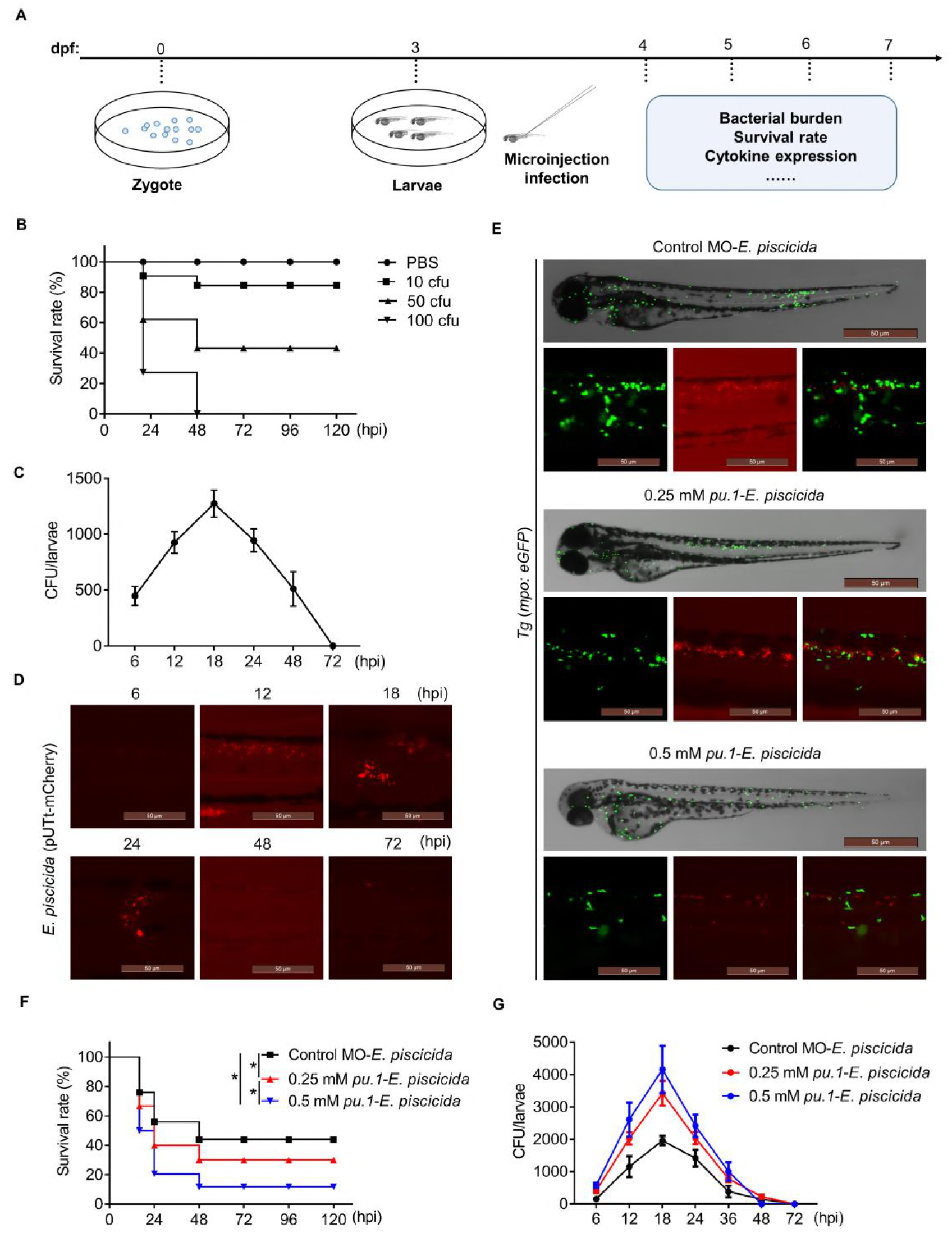
Macrophages and neutrophils are critical for *E. piscicida* infection. (A) Scheme showing the experimental procedure used for the assays of survival rate and bacterial burden. 3-dpf larvae were microinjected with bacteria at the vein tail muscle and survival rate and bacterial burden were determined at the indicated time points. (B) Indicated doses of wild type *E. piscicida* were microinjected into the tail vein muscle of 3-dpf zebrafish larvae, and the survival rate was calculated at indicated days post infection. Experiments used 40 larvae per group, and data shown are from 1 experiment representative of 3 independent experiments. (C) 3-dpf larvae were microinjected with 1 nl suspension of wild type *E. piscicida* (45 cfu/nl) to determine the bacterial colonization in larvae. Data are presented as mean ± SD of 60 larvae per group, 3 independent experiments were analyzed. (D) *In vivo* imaging of the bacterial loading at the tail vein muscle. 3-dpf larvae were microinjected with 1 nl mCherry-labeled wild type *E. piscicida* (45 cfu/nl) at the tail vein muscle. Scale bar=50 μm. (E-G) One-cell stage *Tg* (*mpo:eGFP^+/+^*) embryos were microinjected with 1nl 0.25 mM *pu.1* morpholino to knockdown macrophages, or 1 nl 0.5 mM *pu.1* morpholino to knockdown macrophages and neutrophils, respectively, the control group was microinjected with standard control morpholino. Three days later, the larvae were microinjected with wild type *E. piscicida,* and the bacterial loading was analyzed by fluorescence microscope (E), and the survival rate (F) and bacterial burden (G) were monitored at indicated time points. Images are representative of 3 independent experiments, data are presented as mean ± SD, and the differences in fish survival are analyzed by log-rank (Mantel-Cox) test. * *p*<0.05.

Neutrophils and macrophages are highly motile phagocytic cells that are typically the first responders recruited to sites of tissue infection (25). Previous study has established the model to treat zebrafish larvae with 0.25 mM *pu.1* morpholino, a transcription factor essential for development of myeloid cells, which can suppress the macrophages development, while 0.5 mM *pu.1* morpholino treatment can suppress both macrophages and neutrophils development (26). In this study, in order to characterize further the nature of myeloid cell interaction with *E. piscicida*, we first microinjected 0.25 mM *pu.1* morpholino into embryos, to inhibit the formation of macrophages in *Tg* (*mpo:eGFP*^+/+^) zebrafish larvae (Fig. 1E). Interestingly, we found a comparatively higher mortality in *pu.1* knockdown zebrafish larva following infection with *E. piscicida* (Fig. 1F). Moreover, the bacterial burdens were also comparatively enhanced in 0.25 mM *pu.1* knockdown zebrafish larva (Fig. 1G), which suggesting that macrophages are important for the prevention of *E. piscicida* proliferation. Moreover, we microinjected 0.5 mM *pu.1* morpholino into embryos, to inhibit the formation of both macrophages and neutrophils in *Tg* (*mpo:eGFP*^+/+^) zebrafish larvae, then infected with *E. piscicida*, we found a significantly higher mortality and bacteria burden in both macrophages and neutrophils depletion larva (Fig. 1E), strongly suggests that both macrophages and neutrophils are essential for the prevention of *E. piscicida* proliferation.

### EvpP inhibits neutrophils recruitment to promote *E. piscicida* infection*in vivo*

To characterize the effects of EvpP on the virulence of *E. piscicida*, we microinjected the LD_50_ dose of wild type, Δ*evpP*, and *evpP-*complemented (Δ*evpP*::p*evpP*) *E. piscicida* strains *in vivo*, and monitored the survival and pathogen loads, respectively. We found that the *evpP* mutant strain showed significant attenuation (*p*=0.0248), and the virulence was restored to the same magnitude as that of the wild type in the *evpP-*complemented strain (Fig. 2A). Consistent with the attenuated virulence observed in Δ*evpP* strain, the bacterial burdens were significantly reduced in the infected larvae (Fig. 2B). Interestingly, by utilizing the tail muscle microinjection infection model in *Tg* (*mpo:eGFP*^+/+^) zebrafish larvae with the indicated mCherry-labeled *E. piscicida* strains to establish a direct methods to analysis of leukocyte response to a compartmentalized infection (27, 28), we found that neutrophils were significantly recruited to the Δ*evpP E. piscicida* infection sites, compared with the wild type or *evpP*-complemented *E. piscicida* infection groups (Fig. 2C and D), which suggest that the bacterial T6SS effector EvpP was critical for inhibiting neutrophils recruitment during infection in vivo.

**FIG 2.**
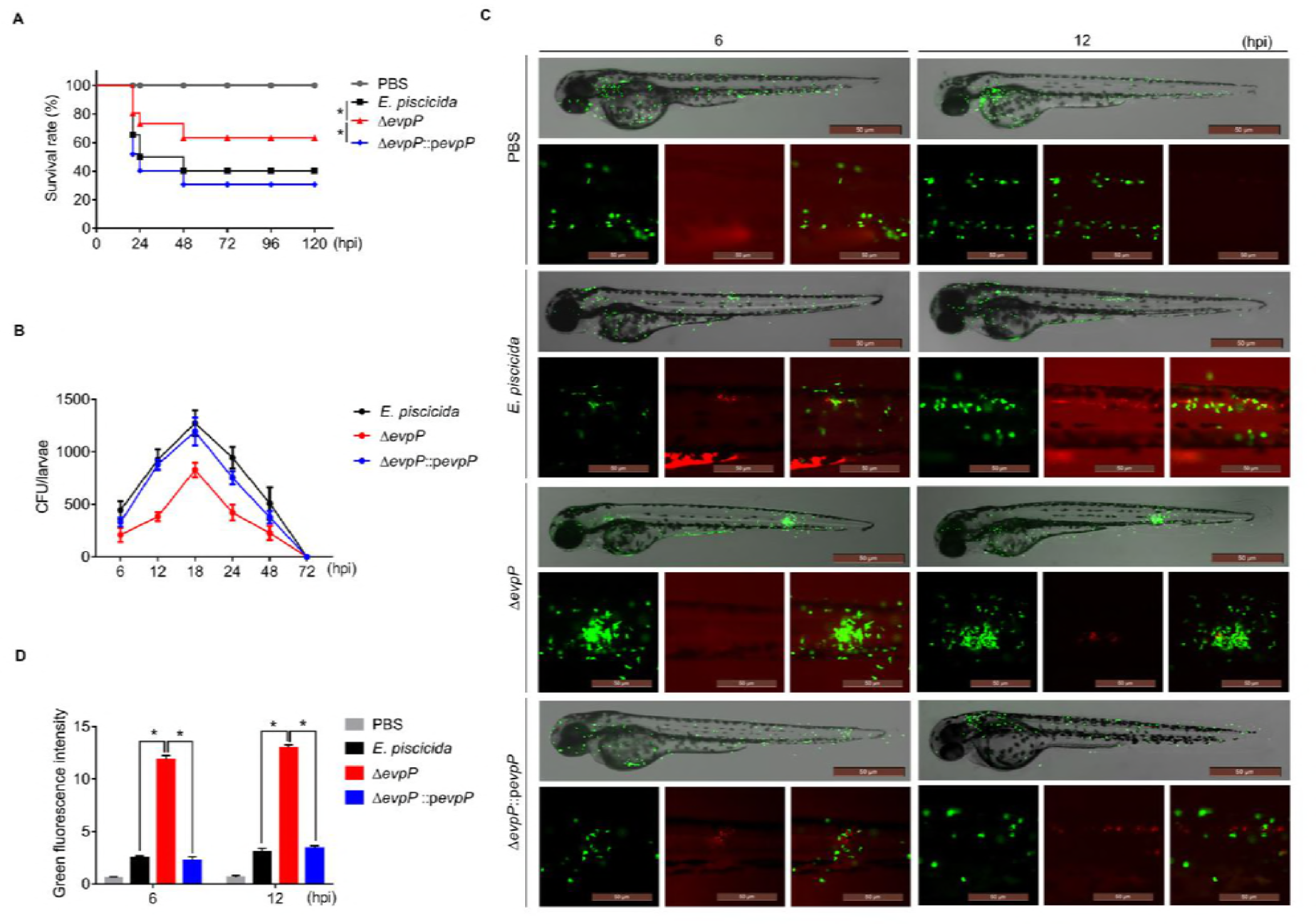
EvpP inhibits neutrophils recruitment to promote *E. piscicida* infection *in vivo*. 3-dpf larvae were microinjected with mCherry-labeled wild type, Δ*evpP,* or Δ*evpP*::p*evpP E. piscicida* at the tail muscle. Survival rate (A) and bacterial burden (B) were monitored at indicated time points. The recruitment of neutrophils to infection site was analyzed by fluorescence microscope at 6 and 12 hpi (C), and the quantification of neutrophil fluorescence intensity were analyzed (D). Images are representative of 3 independent experiments, data are presented as mean ± SD, the differences in fish survival are analyzed by log-rank (Mantel-Cox) test, * *p*<0.05, and the difference between groups are analyzed by student’s *t* test, * *p*<0.05.

### EvpP inhibits neutrophils recruitment via Jnk-MAPK signaling *in vivo*

It has been reported that EvpP may inhibit the phosphorylation of JNK-MAPK signaling during *E. piscicida* infection in mammalian macrophages (22). First, we confirmed that activation of JNK was enhanced in Δ*evpP*-infected zebrafish fibroblasts (ZF4), compared to cells infected with the wild type or *evpP*-complemented *E. piscicida* strains (Fig. S1A). Furthermore, we examined the effects of EvpP on the regulation of JNK-MAPK during bacterial infection *in vivo*. When we infected zebrafish larvae with the indicated *E. piscicida* strains, consistent with the *in vitro* infection experiments, activation of JNK was enhanced in Δ*evpP*-infected larvae, compared to larvae infected with the wild type or *evpP*-complemented *E. piscicida* (Fig. 3A). These results indicate EvpP-mediated manipulation of JNK-MAPK signaling *in vivo*.

**FIG 3.**
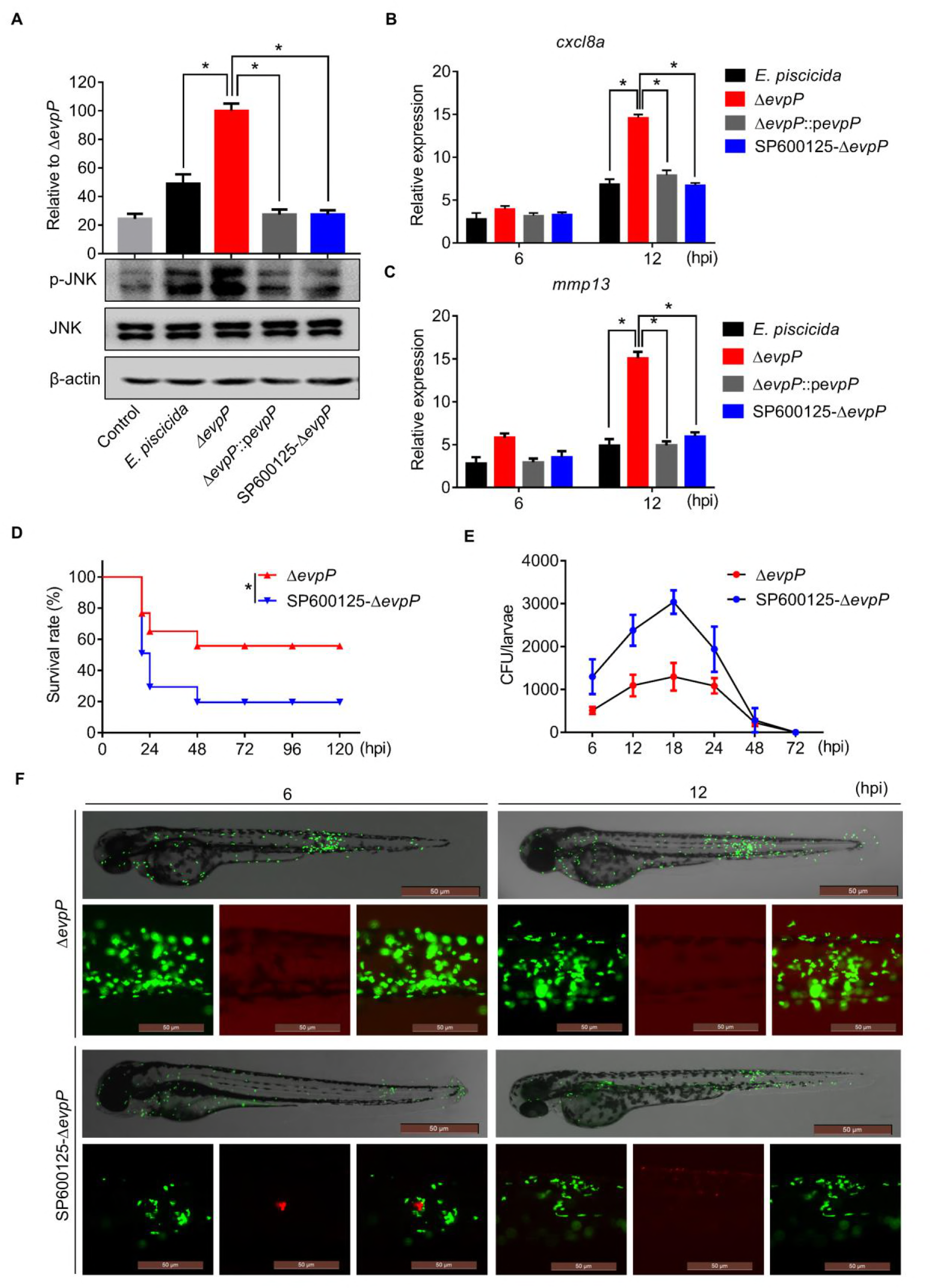
EvpP inhibits neutrophils recruitment via Jnk-MAPK signaling *in vivo*. (A) Quantitative western blot analysis of phosphorylated-JNK (p-JNK) levels of infected-larvae at 12 hpi with or without treatment of JNK inhibitor SP600125 (50 μM). 3-dpf larvae were microinjected with 1 nl PBS, or wild type, Δ*evpP* or Δ*evpP*::p*evpP E. piscicida* (45 cfu/nl); 50 larvae per group were collected for immunoblotting. Results are representative of 3 independent experiments. (B and C) RT-PCR analysis of *cxcl8a* and *mmp13* transcription in infected larvae at indicated timepoints. (D and E) Larva were pretreated with or without 50 μM SP600125, bacterial burden (D) and survival rate (E) of larvae infected with Δ*evpP* were monitored at the indicated timepoints. (F) Larva were pretreated with or without 50 μM SP600125, the neutrophil recruitment to infection site were analyzed by fluorescence microscope. Images are representative of 3 independent experiments, data are presented as mean ± SD, the differences in fish survival are analyzed by log-rank (Mantel-Cox) test, * *p*<0.05, and the difference between groups are analyzed by student’s *t* test, * *p*<0.05.

Since MAPK signaling cascades regulate the transcriptional activation of a wide array of proinflammatory and chemokine genes (29, 30), we further assessed the expression of chemokines in both ZF4 cells and zebrafish larvae infected with wild type and mutant *E. piscicida*. Infection with wild type *E. piscicida* induced the expression of *cxcl8a* and *mmp13* transcripts, which was further enhanced in zebrafish infected with Δ*evpP E. piscicida* (Fig. 3B and C). The enhancement of chemokine expression observed with the EvpP mutant was abolished when ZF4 cells or zebrafish larvae were infected with the *evpP*-complemented *E. piscicida* (Fig. 3B and C). Collectively, these results indicate that EvpP regulates *cxcl8a* and *mmp13* expression by inhibiting JNK-MAPK signaling cascades.

To further assess the role of JNK signaling cascade activation in response to *E. piscicida* infection *in vivo*, we utilized the specific JNK inhibitor SP600125 to pretreat the cells or larvae (30, 31), and found that Δ*evpP*-infection-induced JNK phosphorylation was restored to comparative levels both in infected ZF4 cells and zebrafish larvae groups (Fig. S1A; Fig. 3A), Moreover, SP600125 treatment also inhibited Δ*evpP*-infection enhanced *cxcl8a* and *mmp13* expression(Fig. S1B and C; Fig. 3B and C). Thus, we infected zebrafish with Δ*evpP* with or without SP600125 treatment and monitored the survival and bacterial burden after infection. SP600125 treatment resulted in significant mortality (Fig. 3D), and consistently higher levels of bacterial colonization were detected in SP600125-treated zebrafish larvae (Fig. 3E). Taken together, these results indicate that activation of JNK-MAPK signaling plays a critical role in *E. piscicida* clearance.

In zebrafish, cxcl8a and mmp3 are known to be the most potent chemoattractants, which are responsible for guiding neutrophils through the tissue matrix until they reach sites of injury or infection (31). In this study, we showed that both *cxcl8a* and *mmp13* genes were upregulated in response to Δ*evpP E. piscicida* infection, but the *in vivo* function of EvpP on the migratory behavior of neutrophils during infection remains to be clarified. Here, we found the Δ*evpP E. piscicida* infection-induced recruitment of neutrophils was restored when the zebrafish larvae were pretreated with the specific JNK inhibitor SP600125, which indicates that JNK-MAPK signaling activation plays a critical role in neutrophil immigration (Fig. 3F). Collectively, these results indicate that EvpP plays a critical role in inhibiting the recruitment of neutrophils through the Jnk-MAPK signaling cascade, promoting the bacterial colonization *in vivo*.

### EvpP inhibits neutrophils recruitment through Jnk-caspy-inflammasome cascades *in vivo*

Recently, we have reported that EvpP could inhibit the phosphorylation of Jnk-MAPK to suppress the inflammasome activation during *E. piscicida* infection in mammalian macrophages (22). Thus, to further clarify the downstream signaling of EvpP-regulated Jnk-MAPK activation *in vivo*, we first analyzed the expression of IL-1β during indicated *E. piscicida* infection. Infection with wild type *E. piscicida* induced the expression of *IL-1β* transcripts, which was further enhanced in zebrafish larvae infected with Δ*evpP E. piscicida* (Fig. 4A). while the enhancement of cytokine expression observed with the EvpP mutant was abolished in zebrafish larvae when infected with the *evpP*-complemented *E. piscicida* (Fig. 4A). Furthermore, to analysis the relative caspase-1 activity in zebrafish larvae infected with indicated *E. piscicida* strains, consistently, we found a comparatively increased activation of caspase-1 in zebrafish larvae infected with Δ*evpP E. piscicida* (Fig. 4B). However, when we pretreated the larva with a specific JNK inhibitor SP600125, we found the Δ*evpP E. piscicida* infection-induced *IL-1β* expression and caspase-1 activity was impaired (Fig. 4A and B), which indicates that EvpP-regulated Jnk-MAPK signaling activation plays a critical role in regulating the downstream inflammasome activation *in vivo*.

**FIG 4.**
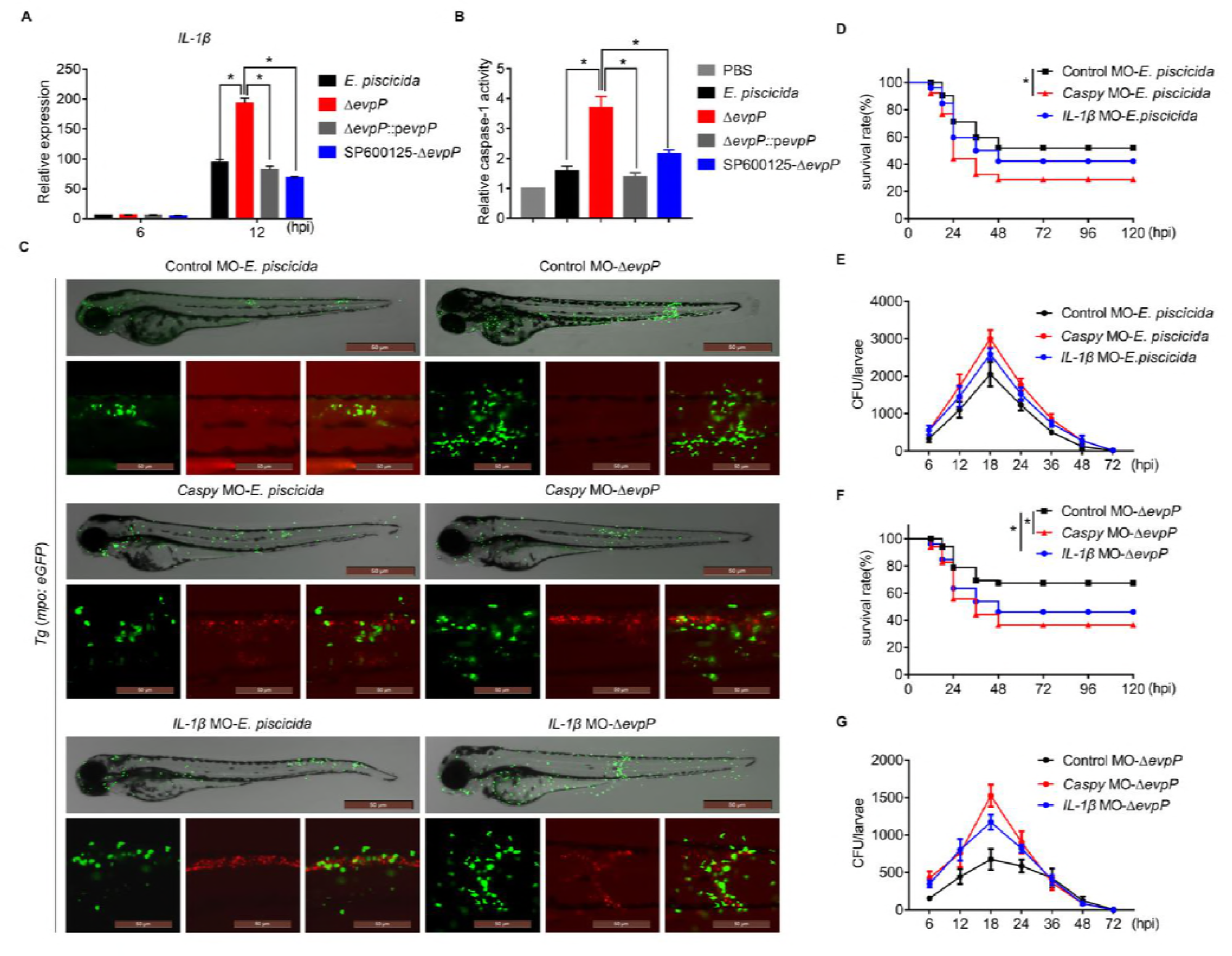
EvpP inhibits neutrophils recruitment through Jnk-caspy-inflammasome cascades *in vivo*. (A) RT-PCR analysis of *IL-1β* transcription in indicated *E. piscicida* infected larvae at indicated timepoints. (B) Relative caspase-1 activity in indicated *E. piscicida* infected larvae was measured by incubating larva homogenate with fluorogenic and chromogenic substrates of caspase-1 (YVAD). (C) One-cell stage *Tg* (*mpo:eGFP^+/+^*) embryos were microinjected with 1 nl 0.75 mM *caspy* morpholino to knockdown caspy, or 1 nl 0.5 mM *IL-1β* morpholino to knockdown IL-1β, and the control group was microinjected with standard control morpholino. Three days later, larvae were microinjected with mCherry-labeled wild type, or Δ*evpP E. piscicida*, the neutrophil recruitment to infection site were analyzed by fluorescence microscope. (D and E) Enumeration of survival rate (D) and bacterial burden (E) of larvae infected with wild type *E. piscicida* were monitored at indicated timepoints. (F and G) Enumeration of survival rate (F) and bacterial burden (G) of larvae infected with Δ*evpP E. piscicida* were monitored at indicated timepoints. Images are representative of 3 independent experiments, data are presented as mean ± SD, the differences in fish survival are analyzed by log-rank (Mantel-Cox) test, * *p*<0.05, and the difference between groups are analyzed by student’s *t* test, * *p*<0.05.

To further analysis the role of inflammasome in regulating neutrophils recruitment, specific caspy (the caspase-1 homolog) or *IL-1β* morpholino was microinjected into *Tg* (*mpo:eGFP*^+/+^) zebrafish larvae embryo, then infected with indicated *E. piscicida* strains. We found the Δ*evpP E. piscicida* infection-induced recruitment of neutrophils was comparatively decreased in both *caspy*‐ and *IL-1β*-MO zebrafish larvae (Fig. 4C), which indicates that the inflammasome signaling activation contribute to the regulation of neutrophil immigration. Moreover, we found a significantly higher mortality in either *caspy*‐ or *IL-1β*-MO zebrafish larvae, and consistently, the bacterial burdens were also comparatively enhanced in either *caspy*‐ or *IL-1β*-MO zebrafish larvae (Fig. 4D-G). Taken together, these results suggest that EvpP could regulate zebrafish larvae Jnk-caspy inflammasome pathways activation, which not only plays critical role in inhibiting neutrophil recruitment, but also highlights the essential role of myeloid cells in response to *E. piscicida* infection *in vivo*.

## DISCUSSION

The progression of infectious disease is determined by dynamic and complex interactions between host defense systems and pathogen virulence factors (32). Pathogenic bacteria encode protein secretion systems that promote invasion of the host’s immune response, thwarting of microbial competitors, and ultimately survival within the host (32). There has been increasing use of zebrafish larvae to study infectious disease, as their optical accessibility and potential for genetic manipulation allows for the visualization of the immune response to infection inside a living intact vertebrate host. For example, injection of *Salmonella enterica serovar* Typhimurium into zebrafish has been key for the discovery of novel concepts in cellular immunity, immunometabolism, and emergency granulopoiesis (33, 34). Recent work has established zebrafish as a model for foodborne enterohaemorrhagic *E. coli* (EHEC) infection, a major cause of diarrheal illness in humans (35). Using the protozoan *Paramecium caudatum* as a vehicle for EHEC delivery, research has shown that zebrafish larvae can be used to study the hallmarks of human EHEC infection, including EHEC-phagocyte interactions in the gut and bacterial transmission to naive hosts (36). In the case of *Shigella*, caudal vein infection of zebrafish was first developed to study *Shigella*-phagocyte interactions and bacterial autophagy *in vivo* (36). In a previous study, we uncovered the mechanism underlying the inhibition by EvpP of Jnk-MAPK pathways, which results in the inhibition of NLRP3 inflammasome activation in mammalian macrophages, thus providing critical insight into this delicate interaction (22). Moreover, during *E. piscicida* infection, it can replicate in a specific vesicle to activate pyroptosis and releases them to the extracellular space (37), but the mechanism of this bacteria arms its virulence effector to resistant the host immune responses *in vivo* remain unknown. Furthermore, intracellular pathogens that replicate within macrophages, such as *S. typhimurium*, must effectively evade pyroptosis in order to stay within an infected cell. Otherwise, detection by the inflammasome activates caspase-1 and triggers pyroptosis, which releases the pathogen to the extracellular space (38), however, the mechanism of bacterial resistance to neutrophil killing remains unknown. Thus, it is necessary to take these advantages to clarify the complexity between macrophages and neutrophils that is regulated by bacterial virulence effectors, and further clarify programmed cell death induced by bacterial infection that shapes innate immune responses *in vivo*. In this study, our results advance the understanding of bacterial T6SS effector EvpP in inhibiting the neutrophil recruitment through Jnk-caspy inflammasome cascades activation *in vivo* (Fig. 5).

**FIG 5.**
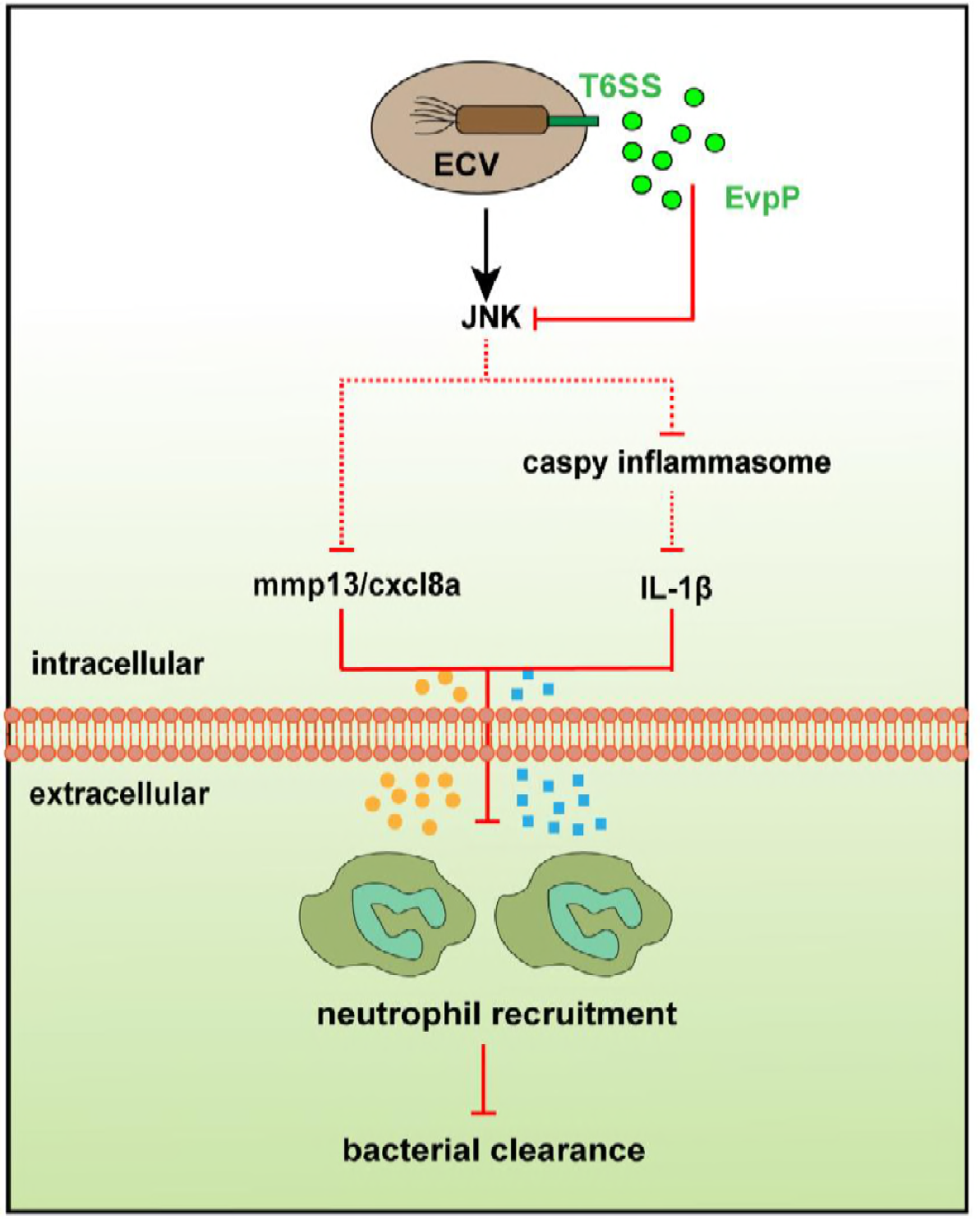
Summary of the proposed mechanism of EvpP *in vivo*. During *E. piscicida* replicates in the *E. piscicida*-containing vacuole (ECV) when it enters the cells, the T6SS effector EvpP could inhibit the phosphorylation of Jnk-MAPK signals, results into the reduced expression of *cxcl8a* and *mmp13*, which contributes to the neutrophil recruitment. Meanwhile, the EvpP inhibited Jnk-MAPK signaling could also inhibit the caspy-inflammasome and IL-1β expression to suppress the neutrophil recruitment, which promotes the colonization of *E. piscicida* in zebrafish. The proposed model suggests a critical role for the *E. piscicida* T6SS effector in modulating the function of immune cells and promoting pathogenesis *in vivo*.

Neutrophils are the most abundant cellular component of the host immune system and are a primary constituent of the innate immune response to invading microorganisms (25). Neutrophil recruitment to areas of inflammation is considered to be the result of the concerted action of several chemoattractants, including not only cytokines, such as IL-1β, but also chemokines, such as cxcl8a (also known as IL-8) or enzymes such as mmp13 (30, 31). These molecules are consistently among the first signals to be expressed and released by the various cell types involved in inflammation (39). Since the genetic tools in zebrafish allow for the generation of transgenic lines with fluorescently labeled cell populations, including neutrophils and macrophages, recent advances in the use of zebrafish to study neutrophils during host-pathogen interactions have been used for mammalian-derived bacteria (31, 40). In addition, the roles of cxcl8 and mmp13 in neutrophil response to tissue injury have been clarified through studies in zebrafish larvae (30, 31). However, it is critical to clarify the importance of neutrophils in host defense against bacterial infection. Here, our results not only confirmed an important function of both macrophages and neutrophils in response to *E. piscicida* infection, but also bifurcates the model to analysis the role of inflammasome activation in regulating neutrophils recruitment during infection, and directly linked the zebrafish caspy-inflammasome activation and neutrophil killing once the pathogen is exposed.

Taken together, this study reveals the first functional characterization of neutrophil migration by a pathogenic T6SS effector in zebrafish larvae, which will shed light on further analysis of the complex and contextual role of bacterial T6SS effectors in modulating the function of macrophages and neutrophils *in vivo*, and offers new insights into the warfare between bacterial weapons and host innate immunological surveillance.

## MATERIALS AND METHODS

### Strains and media

*E. piscicida* strains are listed in Table S1, and were grown at 30°C in tryptic soy broth (TSB) supplemented with antibiotics as appropriate at the following concentrations: colistin (Col), 16.7 μg/ml; ampicillin (Amp), 100 μg/ml.

### Zebrafish strains and maintenance

Zebrafish were obtained from the China Zebrafish Resource Center (CZRC; Wuhan, China). The *Tg* (*mpo:eGFP^+/+^*) line has been previously described (41). Embryos were incubated in E3 medium (5 mM NaCl, 0.17 mM KCl, 0.33 mM CaCl_2_, and 0.33 mM MgSO_4_) containing 0.3 μg/ml methylene blue at 28°C. Experiments were conducted according to protocols approved by the Animal Care Committee, East China University of Science and Technology, Shanghai, China (No. 2006272).

### Infection of *E. piscicida* by microinjection into zebrafish larvae

Bacteria were grown on TSB plates overnight at 30°C, and single colonies of each strain were inoculated into 5 ml TSB supplemented with appropriate antibiotics, grown overnight at 30°C with shaking at 200 rpm, subcultured 1:100 in the same medium, and grown for 4 h without shaking at 30°C to log phase. Then, 1 ml of culture was centrifuged at 4,500× *g* for 10 min to pellet bacteria, and the supernatant was discarded. The bacterial pellet was resuspended in 1 ml sterile PBS, the OD_600_ was measured, and the suspension was diluted to the appropriate concentrations.

Three-dpf zebrafish larvae were mechanically dechorionated and anaesthetized by immersion in 0.02% w/v buffered tricaine (MS-222, Sigma-Aldrich, St. Louis, MO). Embryos were embedded in 2.5% w/v agarose plates and injected individually using pulled glass microcapillary pipettes filled with the appropriate dilution of bacterial suspension. Aliquots of 1 nl` of bacterial suspension were microinjected into tail vein muscle. Injections were performed using pulled borosilicate glass microcapillary injection needles (Sutter Health, Sacromento, CA) and a Milli-Pulse Pressure Injector (Applied Scientific Instrumentation, Eugene, OR). After injection, larvae were placed in petri dishes with E3 medium. The infected larvae were analyzed to detect survival rate, bacterial burden, phosphorylation of JNK *in vivo,* and transcription of cytokines.

For treatment with inhibitors, zebrafish larvae were preincubated 1 h before infection with 50 μM SP600125 (Selleck Chemicals, Boston, MA) in E3 medium, or DMSO (Life Technologies, Carlsbad, CA) as a control. The embryos were maintained in this solution after fin transection over the entire course of the experiment.

For larvae mortality during infection, each indicated bacterial strain was injected into 60 embryos. Following infection, larvae were observed every 24 h, up to 120 hpi, dead embryos were removed, and numbers were recorded at each time point.

For the bacterial burden analysis in infected larvae, 5 zebrafish larvae for each bacterial strain group were transferred individually into a sterile 1.5 ml tube containing 200 μl lysis buffer (1% Triton X-100, Sangon Biotech Co. Ltd., Shanghai, China) and mechanically homogenized on ice. The homogenates were serially diluted and plated onto solid TSB medium to count the bacterial numbers within infected larvae.

### ZF4 cell cultures and infection assays

ZF4 cells (ATCC CRL-2050™, CZRC), established from 1-day-old zebrafish embryos, were cultured in growth medium (GM) consisting of DMEM/F12 (Life Technologies) supplemented with 10% fetal bovine serum (FBS) and seeded in flat bottom 24-well plates (Corning Inc., Corning, NY) at a density of 2 × 10^5^ cells per well and cultured overnight. Before infection, the culture medium was changed to serum-free DMEM/F12 medium (SFM) for 12–16 h. ZF4 was infected at a multiplicity of infection (MOI) of 50, and the bacteria were centrifuged onto cells at 600× *g* for 10 min. For pharmacological pretreatment, the cells were preincubated with SP600125 1 h before infection. The infected cells were detected by immunoblotting and RT-PCR.

### Western blot analysis

Fifty 12-hpi larvae per group were transferred to a 1.5 ml tube. Larvae were homogenized in lysis buffer (20 mM Tris-HCl (pH 7.4), 150 mM NaCl, 5 mM EDTA, 10% glycerol, and 0.1% Triton X-100) containing protease inhibitor cocktail and phosphatase inhibitor (Roche Applied Science, Penzberg, Germany). Protein lysates were obtained by organic solvent precipitation method, protein precipitate was mixed with protein loading buffer, boiled for 10 min and centrifuged (12,000 rpm, 5 min). Protein lysates (10 μl) were separated by SDS-PAGE and transferred to a PVDF membrane (Millipore Sigma, Burlington, MA). The membranes were blocked in 5% w/v nonfat dry milk in TBST. Signals were detected with mouse anti-actin antibody (Ab) (1/5000, Sigma-Aldrich), rabbit anti-phospho-JNK kinases (1/1,000, Cell Signaling Technology, Danvers, MA), and rabbit anti-JNK (1/1,000, Cell Signaling Technology) overnight at 4°C, followed by incubation with the appropriate secondary HRP-conjugated-anti-rabbit Abs (1/2,000, Beyotime Biotechnology, Shanghai, China) and detection with ECL (Cell Signaling Technology). The signal intensities were quantitatively analyzed using NIH ImageJ.

### Quantitative Real-Time PCR analysis

The expression of *cxcl8a* and *mmp13* was evaluated by quantitative real-time RT-PCR of zebrafish larvae and ZF4 cells. Ten infected larvae of each group were sampled at 6 hpi and 12 hpi, and total RNA was isolated using the TRIzol reagent (Life Technologies), treated with DNase I (Promega, Madison, WI) to digest residual genomic DNA, and reverse transcribed using the PrimeScript RT reagent kit (TaKaRa Bio Inc., Kusatsu, Japan). Quantitative RT-PCR was performed using an ABI 7500 Real-Time PCR System (Applied Biosystems, Foster City, CA). Expression of each gene was expressed as the fold change relative to the expression in the PBS control group.

### Microscopic analysis

To observe recruitment of neutrophils and colonization of bacteria, 3-dpf *Tg*(*mpo:eGFP^+/+^*) zebrafish larvae were injected with 1 nl fluorescent bacterial suspension (1 nl of bacterial suspension containing 200 cfu/nl) in the tail vein muscle. At 6 hpi and 12 hpi, images were acquired by a Leica DMI3000B inverted fluorescence microscope (Leica Camera AG, Wetzlar, Germany) to observe the recruitment of neutrophils to the infection site and the colonization of bacteria.

### Microinjection of morpholino nucleotides into zebrafish zygotes

One-cell stage *Tg*(*mpo:eGFP*^+/+^) zebrafish zygotes were microinjected with 1 nl morpholino (Gene Tools, LLC, Philomath, OR) in yolk sac. Accordingly, 0.25 mM *pu.1* morpholino for knockdown of macrophages, 0.5 mM *pu.1* morpholino for full knockdown of macrophages and neutrophils (42), 0.75 mM *caspy* morpholino for caspy knockdown (43), 1 mM *IL-1β* morpholino for IL-1β knockdown (44). Green fluorescence was observed to confirm the knockdown of neutrophils. Morpholino-treated larvae were then infected as described above.

### Caspase-1 activity assays

The caspase-1 activity was determined with the fluorometric substrate Z-YVAD 7-Amino-4-trifluoromethylcoumarin (Z-YVAD-AFC, caspase-1 substrate VI, calbiochem) as destribed preciously (45). In brief, 25-35 larvae were lysed in hypotonic cell lysis buffer (25 mM 4-(2-hydroxyethyl) piperazine-1-ethanesulfonic acid, 5 mM dithionthreitol, 1:20 protease inhibitor cocktail (Sigma-Aldrich), pH 7.5) on ice for 10 min. For each reaction, 10 μg protein were incubated for 90 min at 23°C with 50 μM YVAD-AFC and 50 μl of reaction buffer (0.2% 3-[(3-cholamidopropyl) dimethylammonio]-1-propanesulfonate (CHAPS), 0.2 M 4-(2-hydroxyethyl) piperazine-1-ethanesulfonic acid, 20% sucrose, 29 mM dithiothreitol, pH 7.5). After the incubation, the fluorescence of the AFC released from the Z-YVAD-AFC substrate was measured using a SpectraMax M5 fluorescent plate reader (Molecular Devices) with an excitation wavelength of 405 nm and an emission wavelength of 492 nm.

### Statistical Analysis

Statistical analysis was performed using Graphpad Prism (GraphPad Software Inc., La Jolla, CA). All data are representative of at least 3 independent experiments and are presented as the mean ± standard deviation (SD). Differences between 2 groups were evaluated using Student’s *t* test. One-way ANOVA test was used to analyze differences among multiple groups. Differences in fish survival were assessed using the log-rank (Mantel-Cox) test. Statistical significance was defined as * *p*<0.05.

## ACKNOWLEDGEMENTS

This work was supported by the National Natural Science Foundation of China No. 31472308, 31622059 (Q.L.) and the Fundamental Research Funds for the Central Universities No. 222201714022 (D.Y.). Dahai Yang was supported by the Young Elite Scientists Sponsorship Program by CAST No. 2016QNRC001, Shanghai Pujiang Program No.16PJD020, Shanghai Chenguang Program No.16CG33, and the Talent Program of the School of Biotechnology of the East China University of Science and Technology, Shanghai, China.

## AUTHOR CONTRIBUTIONS

Q.L., and D.Y. conceived the study; J.T. conducted the majority of experiments with help from X.Z. and Z.W.; Y.Z. provided expert advice and critical review of the manuscript. D.Y., Q.L. and J.T. wrote the manuscript; all authors discussed the results and commented on the manuscript.

## ORCID

Dahai Yang, http://orcid.org/0000-0001-6602-8653

Qin Liu, https://orcid.org/0000-0002-5465-1189

## Supporting information

**FIG S1.**
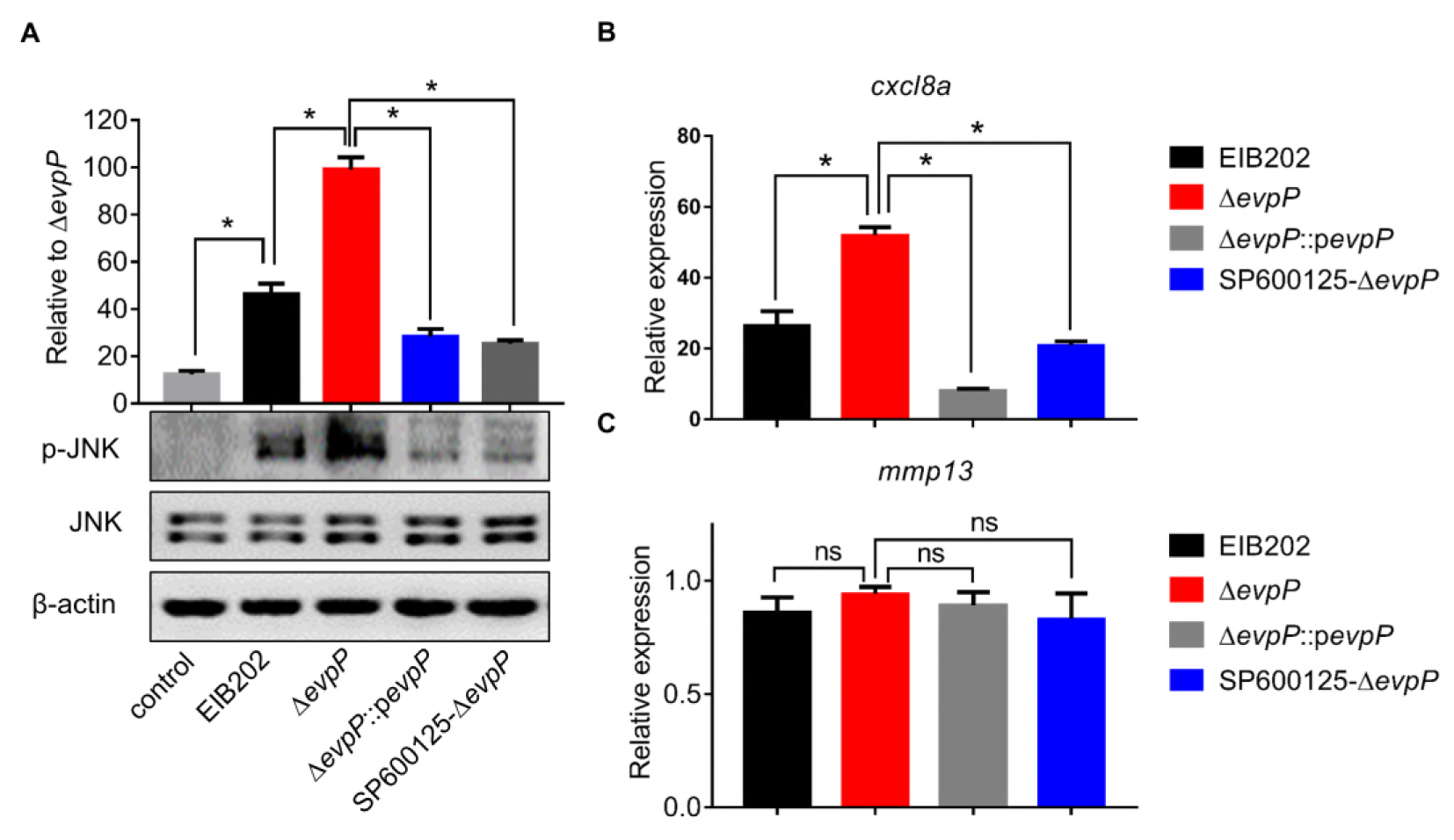
Role of EvpP in regulating JNK-MAPK signaling cascades in infected ZF4 cells. ZF4 cells were pretreated with or without DMSO or JNK inhibitor (SP600125, 40 μM, 1 h) and then infected with or without wild type, Δ*evpP*, or Δ*evpP*::p*evpP E. piscicida* for 3 h at an MOI of 50. The phosphorylation of JNK (P-JNK) was analyzed by immunoblotting, and the transcription of *cxcl8a* and *mmp13* was analyzed by RT-PCR. (A) The phosphorylation of JNK was analyzed by immunoblotting. (B and C) RT-PCR analysis of *cxcl8a* and *mmp13* transcription in indicated *E. piscicida* infected ZF4 cells. Data are presented as mean ± SD, the difference between groups are analyzed by student’s *t* test, * *p*<0.05; ns = not significant.

**Table S1.** Strains and plasmids used in this study

| Strains and plasmids | Description | References |
| --- | --- | --- |
| *Edwardsiella piscicida* |  |  |
| **EIB202** | Wild type, Col<sup>r</sup>, Cm<sup>r</sup> | CCTCC# M208068 |
| **$\Delta evpP$** | EIB202, in-frame deletion of ETAE_2428 | 26 |
| **$\Delta evpP::pevpP$** | $\Delta evpP$, complemented with pUTt-0456-ETAE_2428 | 25 |
| *Escherichia coli* |  |  |
| **DH5α** | α-Complementation | Invitrogen™ |
| **Plasmids** |  |  |
| **pUTt** | Promoter screening vector, pUC18 derivative with lac promoter and MCS deleted, and rrnBT1T2 terminator and MCS from pBV220 inserted |  |
| **pUTt-p0456-*mCherry*** | pUTt-pBAD containing fragment of ETAE_0456 promoter and *mCherry*, Amp<sup>r</sup> | This study |
| **pUTt-p0456-*evpP*** | pUTt-pBAD containing fragment of ETAE_0456 promoter and *evpP*-HA fusion, Amp<sup>r</sup> | This study |
| **pUTt-p0456-*evpP*-*mCherry*** | pUTt-pBAD containing fragment of ETAE_0456 promoter, *evpP*-HA, and *mCherry*, Amp<sup>r</sup> | This study |

**Table S2.**
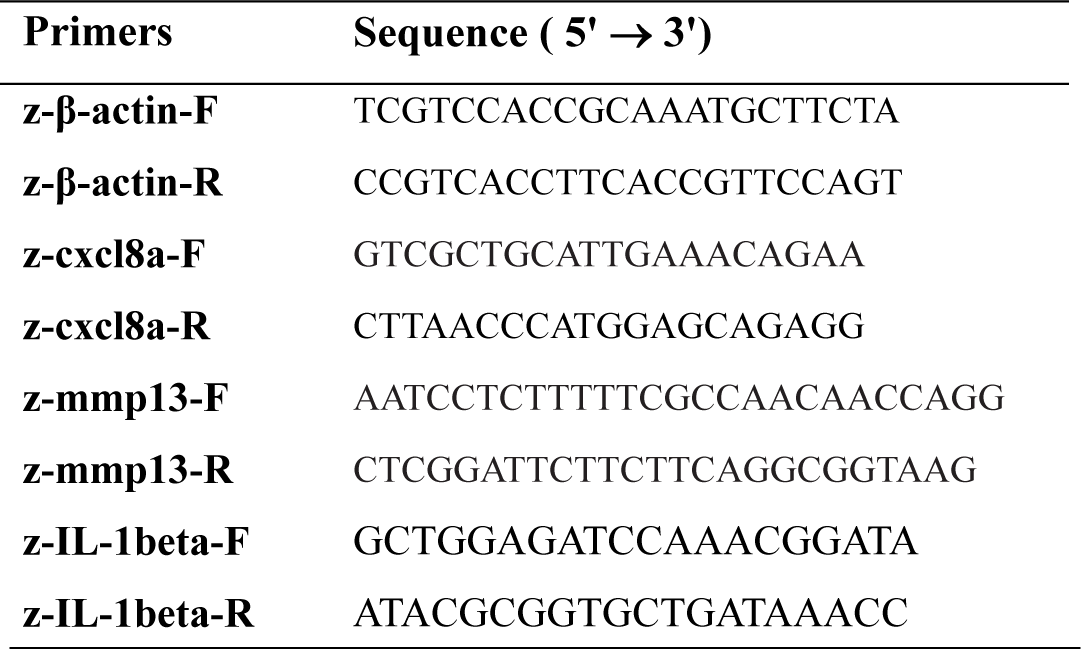
Primers used in this study

